# Chromosomal arrangement of synthetic lethal gene pairs: repulsion or attraction?

**DOI:** 10.1101/2020.05.05.078626

**Authors:** Sayed-Rzgar Hosseini, Bishoy Wadie, Evangelia Petsalaki

**Affiliations:** EMBL-EBI, Wellcome Genome Campus, CB10 1SD, Hinxton, Cambridgeshire, UK; Open Targets, Wellcome Genome Campus, Cambridgeshire, UK; Trinity College Cambridge, University of Cambridge, Cambridge CB2 1TQ, UK

**Keywords:** Synthetic lethality, Genome organization, Yeast, Co-expression, Protein-protein interaction, Paralogs

## Abstract

Synthetic lethal interactions are of paramount importance both in biology and in medicine, and hence increasing efforts have been devoted to their systematic identification. Our previous computational analysis revealed that in prokaryotic species, synthetic lethal genes tend to be further away in chromosomes than random (i.e. repulsion), which was shown to provide bacterial genomes with greater robustness to large-scale DNA deletions. To test the generalizability of this observation in eukaryotic genomes, we leveraged the wealth of experimentally determined synthetic lethal genetic interactions of yeast that are curated in the BioGRID (Biological General Repository for Interaction Datasets) database. We observed an opposite trend that is the genomic proximity of synthetic lethal gene pairs both on the 2D and 3D chromosomal space of the yeast genome (i.e. 2D and 3D attraction). To gain mechanistic insights into the origin of the attraction of synthetic lethal gene pairs in *S. cerevisiae*, we characterized four classes of genes, in which synthetic lethal interactions are enriched and partly explain the observed patterns of genomic attraction: i) gene pairs operating on the same pathways, 2) co-expressed genes, 3) gene pairs whose protein products physically interact and 4) the paralogs. However, our analysis revealed that the contribution of these four types of genes is not sufficient to fully explain the observed 2D and 3D attraction of synthetic lethal gene pairs and hence its evolutionary origin still remains as an open question.

**Significance statement:** Unravelling the organizing principles underlying gene arrangements is one of the fundamental questions of research in evolutionary biology. One understudied aspect of this organization is the relative chromosomal arrangement of synthetic lethal gene pairs. In this study, by analyzing a wealth of synthetic lethality data in yeast, we provide evidence that synthetic lethal gene pairs tend to be attracted to each other both on 2D and 3D chromosomal space of the yeast genome. This observation is in sharp contrast with the repulsion of synthetic lethal (metabolic) gene pairs that we observed previously in bacterial genomes. Characterizing the evolutionary forces underlying this genomic pattern in yeast can open the door towards an evolutionary theory of genome organization in eukaryotes.

## Introduction

Synthetic lethal interactions occur when simultaneous perturbations of two genes cause cellular or organismal death, whilst perturbation of individual genes does not affect viability (Nijman 2011). Their impact on cellular viability implies that synthetic lethal (SL) interactions potentially have been optimized under strong selective pressure in the course of evolution (Dixon et al. 2008). Previously, features of SL gene pairs such as their co-functionality (Srivas et al. 2016), subcellular colocalization (Wong et al. 2004) and conserved patterns of protein-protein interactions (Benstead-Hume et al. 2019) have been investigated to some extent. However, their chromosomal arrangement remains poorly studied, in spite of the well-known functional relevance of the arrangement of genes on prokaryotic (Adams et al. 1992) and eukaryotic (Trowsdale 2002; Hurst et al. 2004) chromosomes.

Previously, we observed in bacterial genomes, the “repulsion of synthetic lethal gene pairs” (Hosseini & Wagner 2018), namely that SL (metabolic) gene pairs tend to be positioned far from each other on bacterial chromosomes, and we provided evidence that this repulsion might be a signature of evolutionary adaptation to pervasive gene deletions (Mira et al. 2001; Kunin & Ouzounis 2003; Nilsson et al. 2005; Lee et al. 2012; Koskiniemi et al. 2012; Albalat & Cañestro 2016) that exert significant selective pressure on bacterial genomes (Sung et al. 2016). If a pair of synthetic lethal genes is closely located on a chromosome (i.e. attracted, Figure 1a), they are likely to be hit simultaneously by a single deletional event and thus cause lethality. Conversely, if they are located far from each other (i.e. repulsed, Figure 1b), only one of the genes is likely to be hit during a deletional event, conferring robustness to such deletional events.

**Figure 1.**
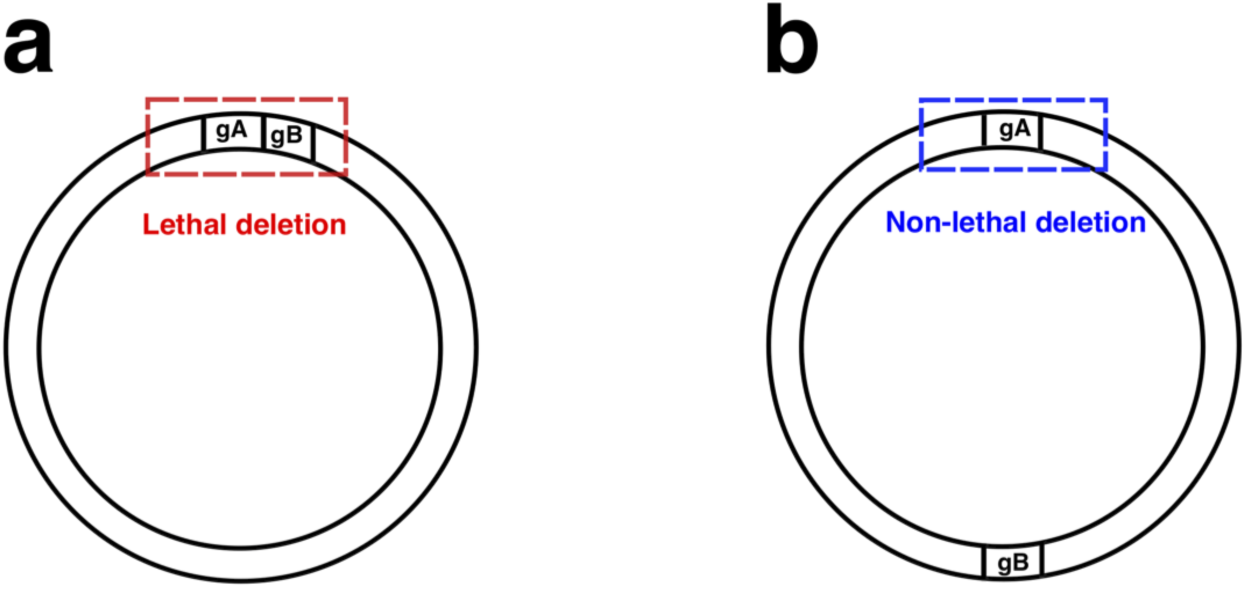
Repulsion of synthetic lethal gene pairs and deletional robustness. The circles represent bacterial chromosomes in which two synthetic lethal genes are in attraction (panel **a**) and repulsion (panel **b**). Deletion of a DNA segment of a given size causes the cell to die when both genes are affected (red rectangle, **a**), but not when only one of the genes is deleted (blue rectangle, **b)**.

However, the importance of the role of large DNA deletions in the evolution of eukaryotic genomes is not comparable to that of prokaryotes. Therefore, we hypothesized that repulsion of synthetic lethal genes would not be observed on eukaryotic chromosomes. Furthermore, eukaryotic genomes are strongly influenced by gene duplication events (Friedman & Hughes 2001; Qian & Zhang 2014), which generate closely located paralog gene pairs accounting potentially for a significant proportion of the synthetic lethal genetic interactions (De Kegel & Ryan 2019; Dede et al. 2020). Moreover, studies on budding yeast as a eukaryotic cell have indicated that gene pairs showing negative genetic interactions tend to participate in closely related biological processes (Kelley & Ideker 2005; Costanzo et al. 2010; Bellay et al. 2011; Costanzo et al. 2016) and hence might be co-expressed. Finally, it was shown (Cohen et al. 2000) that the closely located genes on yeast chromosomes tend to be co-expressed. These observations altogether, raise the possibility of observing an opposite trend that is the enrichment of closely located synthetic lethal gene pairs (i.e. attraction) in eukaryotic chromosomes.

Fortunately, recent high-throughput techniques such as SGA (Synthetic Gene Array (Tong et al. 2001)) or dSLAM (Diploid-based Synthetic Lethality Analysis with Microarrays (Pan et al. 2007)), which in yeast are frequently followed by rigorous classical confirmatory methods such as tetrad dissection, random sporing or spot dilution assays to minimize the false discovery rate, have resulted in thousands of high quality experimentally validated synthetic lethal interactions curated in the BioGRID database (Oughtred et al. 2018). In this study, we leveraged this data to test the above hypothesis in the budding yeast as a well-studied eukaryotic species. Furthermore, the availability of a comprehensive three-dimensional chromosome map in yeast (Duan et al. 2010) encouraged us to extend the analysis to investigate the 3D spatial attraction or repulsion of synthetic lethal gene pairs.

## Materials and methods

### Yeast genomic information

In this study, we focused on *S. cerevisiae* genome, which includes 7926 distinct genes (Supplementary table S1). The whole genome annotation of the yeast was obtained using the Bioconductor R packages, AnnotationDBI based on the corresponding TxDB and org.db (Gentleman et al. 2004; Pages et al. 2018). The chromosomal location and genomic coordinates of genes were retrieved using rentrez R package v1.2.2 (Winter 2017) based on the entrez_id provided in the extracted datasets.

### Synthetic lethality data

Synthetic lethal (SL) gene interactions were obtained from the BioGRID database; release 3.5.184 (Oughtred et al. 2018), which includes 12,552 SL gene pairs for *S. cerevisiae* (supplementary table S2). Note that this is a strictly conservative collection of SL gene pairs in the sense that candidate SL interactions identified by high-throughput methods such as SGA have further been confirmed by classical orthogonal methods such as tetrad dissection or random sporing, which ensure minimum false discovery rate.

### 3D model of the yeast genome

We re-analyzed a previously published three-dimensional genome of yeast (Duan et al. 2010), which has identified chromosomal interactions genome-wide by coupling chromosome conformation capture-on-chip (Simonis et al. 2006) and massively parallel sequencing. The method captures the 3D chromosomal information from the interacting HindIII fragments. Among 8,759,221 potential pairs, 65,683 distinct interacting fragment pairs were identified. We consider a given gene as 3D proximate, if an interacting pair of HindIII fragments are each located within one of the given gene pairs. We identified 188,847 total gene pairs and 206 SL gene pairs as 3D proximate based on this definition. Alternatively, a more moderate criterion would consider a gene pair as 3D proximate, if an interacting pair of HindIII fragments are each located within a given genomic distance (half of the HindIII fragment size) from one of the given gene pairs. Considering the HindIII fragment size as 1 KB, we identified 577,586 total gene pairs including 407 SL pairs among them as 3D proximate (supplementary table S3).

### KEGG pathway analysis

KEGG pathways associated with each gene were retrieved from the KEGG database using the KEGGREST R package (Dan Tenenbaum 2018). We identified 36,071 total gene pairs including 515 SL gene pairs with Jaccard similarity of one, as gene pairs belonging to exactly the same pathways (Supplementary table S4).

### Identification of co-expressed genes

We analyzed the yeast microarray gene expression compendia from SPELL (Serial Pattern of Expression Levels Locator; version 2.0.3) database (Hibbs et al. 2007), which includes genome-wide expression profile from a total of 752 datasets representing 15,475 microarray experiments. To calculate the co-expression of a given gene pair from multiple datasets, we followed the same procedure as in (Hibbs et al. 2007). First, we applied Singular Value Decomposition (SVD) on every dataset separately to minimize the impact of noise in the data. SVD decomposes the original *m* × *n* data matrix, *X*, into three matrices of the form: 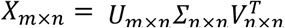 where *U* is the projection of each original data vector in the new basis. Next, for a given dataset, we calculated the Pearson’s correlation coefficient of the expression profiles (in the U matrix) between every gene pair, which was then subjected to Fisher’s z-transformation in order to obtain normally distributed scores for each given dataset. Finally, these quantities were standardized by subtracting the mean and dividing by the standard of the z-scores of the given dataset to get standard normal scores (∼ *N*(*0,1*)) for every dataset. The final score of a given gene pair is quantified as the median of their z-scores among all datasets in which the expression profiles of both genes have been measured. We only considered gene pairs in which both genes were present together in at least 10 datasets. We identified gene pairs as “co-expressed” if their median z-scores >1 (i.e. one standard deviation above the mean), which resulted in a total of 97,815 co-expressed gene pairs including 305 synthetic lethal interactions among them (Supplementary table S5).

### Protein-protein interaction data

We collected the physical protein-protein interaction data from the BioGRID database (Oughtred et al. 2018) resulting in 110,904 total gene pairs, whose protein products physically interact (including 1552 SL gene pairs among them; Supplementary table S6).

### Paralog genes

Paralogous genes were obtained from the PANTHER classification system (Protein ANalysis THrough Evolutionary Relationships; release 15.0) (Mi et al. 2019). UniprotKB identifiers from PANTHER dataset were mapped to entrez_id using “Retrieve/ID mapping” web-based tool from Uniprot (Consortium 2019). We identified 1590 paralog gene pairs including 90 SL gene pairs among them (Supplementary table S7).

### Statistical analyses

We generated the null distribution of genomic distances, by calculating the genomic distance among all gene pairs. To calculate the expected number of synthetic lethal gene pairs located on the same chromosome or in a genomic distance of shorter than a given threshold (e.g. 25 KB), under the null hypothesis, we multiplied the fraction of gene pairs on the same chromosome or within the defined genomic distance in the null distribution by the number of known synthetic lethal gene pairs. Similarly to quantify the expected number of synthetic lethal gene pairs of a given type (e.g. 3D proximate, paralogs, or co-expressed ones, etc.) under the null distribution, we multiplied the fraction of gene pairs of the given type (among all possible gene pairs) by the number of known SL gene pairs in our dataset. Finally, to calculate the statistical significance, we used Chi-squared test. All the statistical analyses and data visualization were done using R software version 3.6.1.

## Results

We examined the distribution of synthetic lethal gene pairs on the genome of *S. cerevisiae*, which has 16 distinct chromosomes. We first observed that 1109/12552 (8.8%) SL gene pairs, compared to 931.32 (7.4%) expected by the null distribution, were located on the same chromosome (Figure 2a). Thus, synthetic lethal gene pairs are to some extent attracted to each other on the inter-chromosomal level in *S. cerevisiae* (*χ*^2^=33.9 and p-value < 10^−8^). Looking into the distribution of the SL gene pairs on the same chromosome, we found that there is significant enrichment of synthetic lethal interactions in shorter genomic distances (Figure 2b and 2c). For example, 161 gene pairs are closely located (genomic distance <25 KB), compared to 63.4 that was expected from the null distribution (*χ*^2^=149.9 and p-value < 10^−33^; Figure 2b and Supplementary table S8). Furthermore, we found that the enrichment of SL gene pairs is statistically significant up to genomic distance of almost 500 KB (Figure 2c). Thus, synthetic lethal gene pairs are attracted to each other on the yeast chromosome (i.e. 2D attraction).

**Figure 2.**
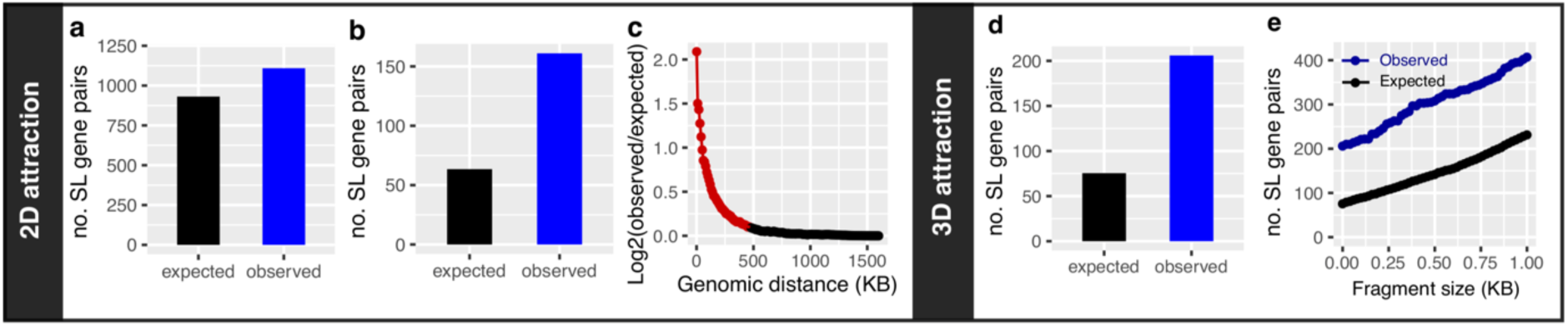
Chromosomal attraction of synthetic lethal gene pairs in the yeast genome (2D and 3D). Vertical axes indicate respectively the number of expected (blue) versus observed (black) **a)** synthetic lethal gene pairs located on the same chromosome, **b)** closely located synthetic lethal gene pairs (with genomic distance<25 KB), **c)** the ratio of the observed versus expected synthetic lethal gene pairs (in log2 scale) located within a given genomic distance (*x*-axis). The red points indicate the genomic distances below which the synthetic lethal gene pairs are significantly enriched (P value of the Chi-square test<0.05), **d)** The expected (balck) versus observed (blue) number of 3D proximate synthetic lethal gene pairs ((Duan et al. 2010); see Methods), and **e)** the expected (blue curve) versus the observed (black curve) number of 3D proximate synthetic lethal gene pairs as a function of HindIII fragment size (*x*-axis).

Next, we examined the attraction or repulsion of synthetic lethal gene pairs on the 3D model of the yeast genome (Duan et al. 2010). To this end, we characterized all gene pairs, which were considered as 3D proximate based on the existence of a pair of 3D interacting HindIII fragments each located within one of the given gene pairs. We observed that 206/12552 (1.6%) SL gene pairs are 3D proximate that is significantly higher than 75.5 (0.6%) expected from the null distribution (Figure 2d; *χ*^2^=225.7 and p-value < 10^−50^). Alternatively, a less strict criterion to consider a given gene pair as 3D proximate, would require an interacting pair of HindIII fragments are each located within a given genomic distance (half of the HindIII fragment size) from one of the given gene pairs. Figure 2e indicates that the enrichment of the 3D proximate SL genes is still preserved in various HindIII fragment sizes. For example, considering the HindIII fragment size as 1 KB, we observed 407/12552 (3.2 %) 3D proximate SL gene pairs that is significantly higher than expected 230.8 (1.8 %) from the null distribution (*χ*^2^=134.4 and p-value < 10^−30^). Therefore, synthetic lethal gene pairs are attracted to each other on the yeast 3D chromosomal space (i.e. 3D attraction).

Our next aim was to characterize the potential factors, which might explain the 2D and 3D attraction of SL gene pairs. For this purpose, we identified four classes of gene pairs (see Methods), in which synthetic lethal interactions are observed significantly more frequently than expected from the null distribution: i) gene pairs whose protein products operate in exactly the same pathways (observed: 515/36071 (1.43%), expected: 14.4 (0.04%), *χ*^2^=17382.2 and p-value < 10^−300^, Figure 3a), ii) co-expressed gene pairs (observed: 305/97815 (0.31%), expected: 39.1 (0.04%), *χ*^2^=1808.3 and p-value < 10^−300^, Figure 3b), iii) gene pairs, whose protein products physically interact (observed: 1552/110904 (1.4%), expected: 44.3 (0.04%), *χ*^2^=51270.7 and p-value < 10^−300^, Figure 3c), and iv) paralogs (observed: 90/1590 (5.66%), expected: 0.6 (0.04%), *χ*^2^=12567.3 and p-value < 10^−300^, Figure 3d).

**Figure 3.**
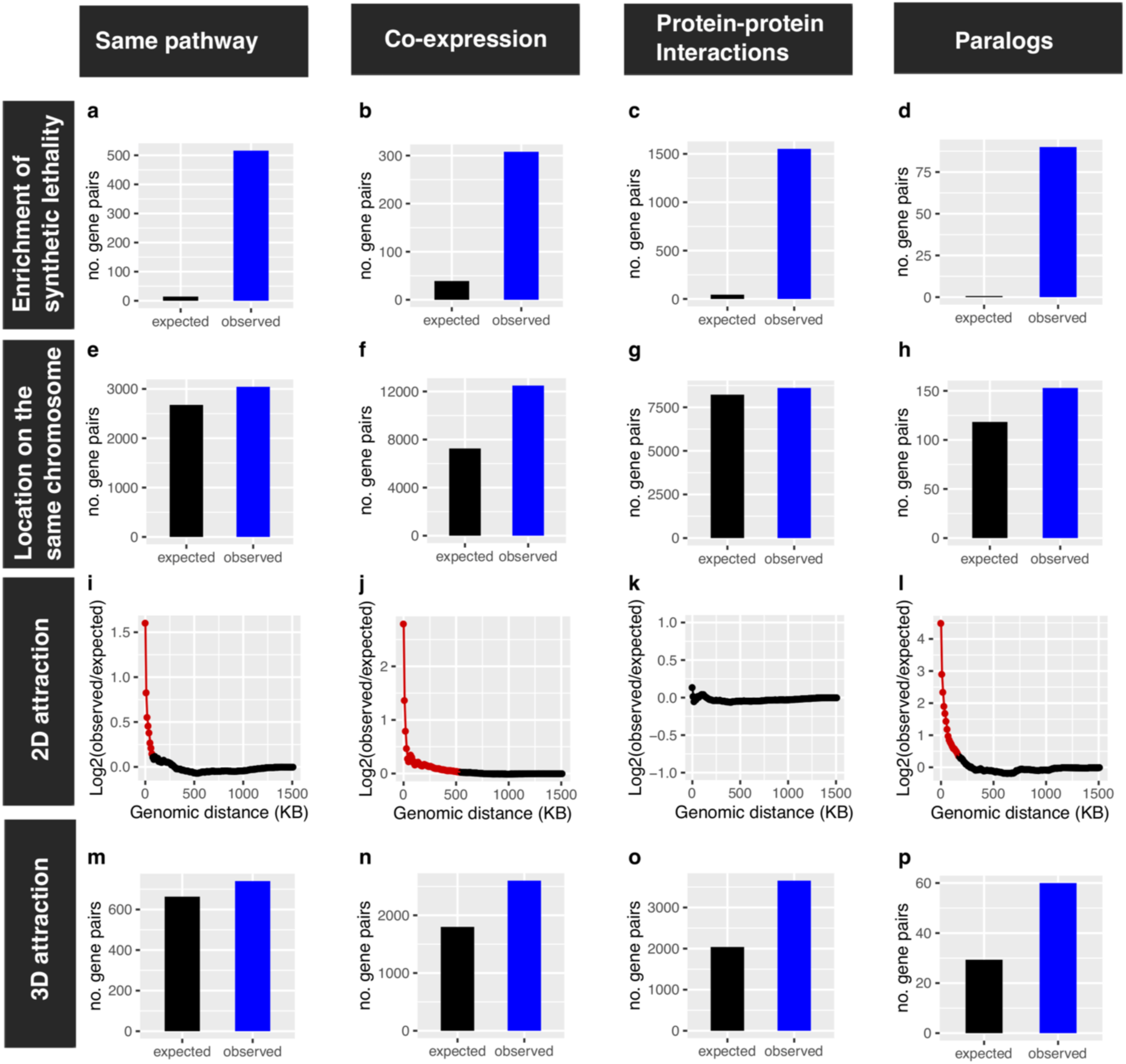
The potential drivers of the attraction of synthetic lethal gene pairs in the yeast genome. The panels in each of the four columns correspond to one of the potential drivers of the attraction of synthetic lethal gene pairs in the *S. cerevisiae* genome. From left to right: i) gene pairs belonging to the same pathway, ii) co-expressed gene pairs, iii) gene pairs, whose protein products physically interact, and iv) paralog gene pairs. Panels in the first row **(a-d)**, show the expected (black) versus the observed (blue) number of synthetic lethal gene pairs. The panels of the second row **(e-h)** illustrate the expected (black) versus observed (blue) number of gene pairs, which are located on the same chromosome. Panels in the third row **(i-l)** show the enrichment of closely located gene pairs (i.e. 2D attraction). The vertical axes indicate the ratio of the observed versus expected gene pairs (in log2 scale) located within a given genomic distance (*x*-axis). The red points highlight the genomic distances in which the given type of gene pairs is significantly enriched (P value of the Chi-square test<0.05). Finally, the panels in the fourth row **(m-p)** illustrate observed (blue) versus expected (black) number of gene pairs in chromosomal regions, which interact in the 3-dimensional chromosomal space of the budding yeast (considering the HindIII fragment size as one KB).

Then, we examined the distribution of these 4 classes of gene pairs over the yeast 2D and 3D genome. We observed that all of them tend to be located on the same chromosome more frequently than expected by chance (Figures 3e-h). Moreover, three classes of gene pairs (all expect those participating in PPI interactions) are enriched among closely located genes, and hence have contributed to the 2D attraction of SL gene pairs (Figures 3i-l). Furthermore, we observed that all four types of gene pairs are significantly enriched among the 3D proximate genes and hence have contributed to the 3D attraction of SL gene pairs as well: class i) observed: 740/36071 (2.05%), expected: 663.4 (1.84%), *χ*^2^=8.8 and p-value =0.003 (Figure 3m), class ii) observed: 2600/97815 (2.65%), expected: 1798.8 (1.84%), *χ*^2^=358.8 and p-value < 10^−78^ (Figure 3n), class iii) observed: 3654/110904 (3.3%), expected: 2039.6 (1.84%), *χ*^2^=1277.9 and p-value < 10^−280^ (Figure 3o), and class iv) observed: 60/1590 (3.77%), expected: 29.2 (1.84%), *χ*^2^=32.35 and p-value < 10^−7^ (Figure 3p).

We then checked the overlap between these four classes to see how independent they are. We observed that the majority of the gene pairs are specific to one of the given classes and only a minor fraction of them is shared across the classes (Figures 4a and 4c). However, when we focus exclusively on synthetic lethal gene pairs, we observe noticeable overlap particularly among the first three classes (Figures 4b and 4d). The protein products of 43.8% of the same-pathway SL gene pairs (class I) and 33.3% of co-expressed SL gene pairs (class II) also interact physically (shared with class III), which underscores the importance of protein-protein interaction in the enrichment of synthetic lethal interactions among the same-pathway and co-expressed gene pairs.

**Figure 4.**
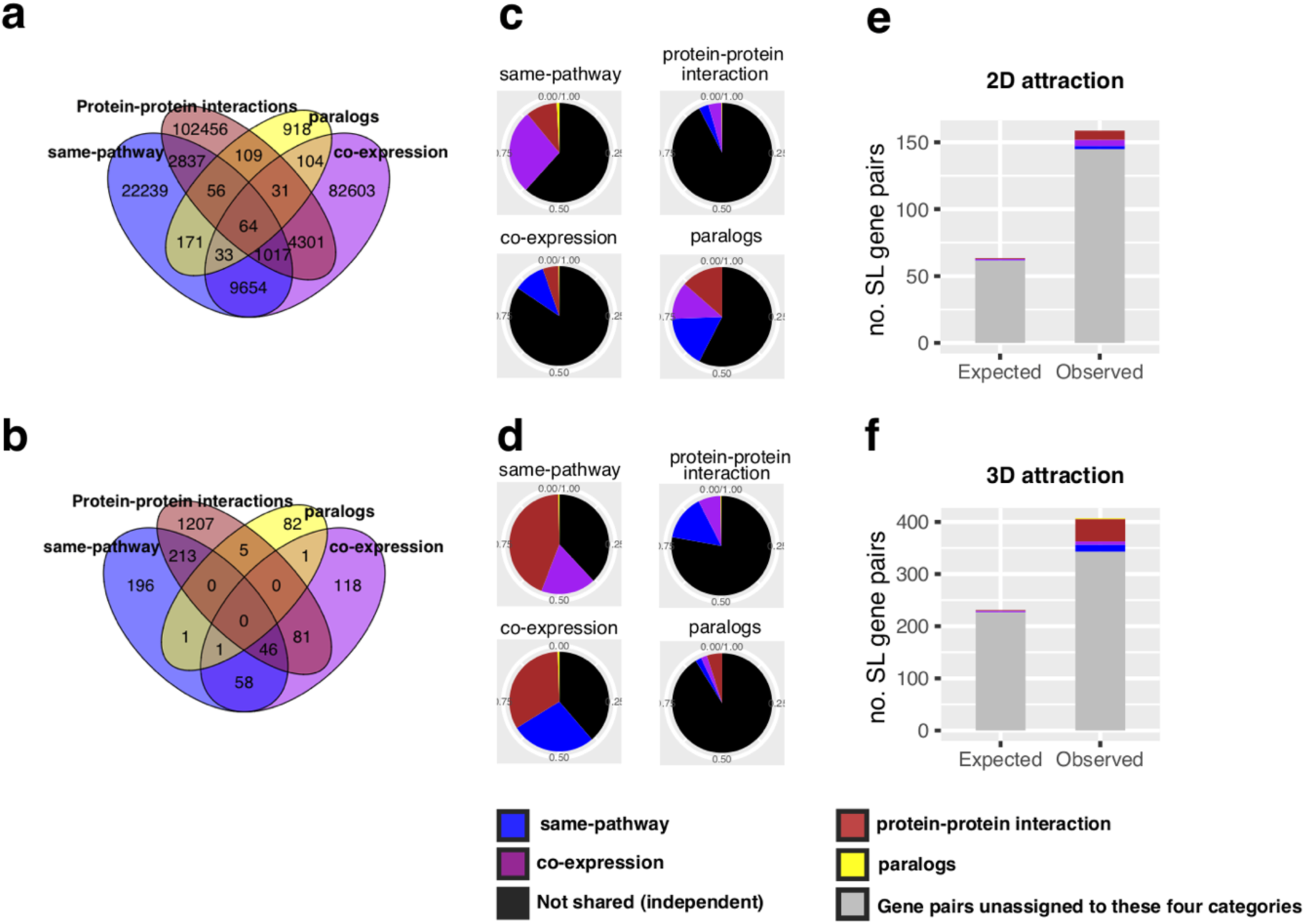
Independence and insufficiency of the four potential drivers of the attraction of synthetic lethal gene pairs in the yeast genome. Venn diagrams (panels **a** and **b**) indicate the overlap between the four types of gene pairs i) gene pairs belonging to the same pathway (blue), ii) co-expressed gene pairs (purple), iii) gene pairs, whose protein products physically interact (brown), and iv) paralog gene pairs (yellow), both among all gene pairs (panel **a**) and specifically among synthetic lethal ones (panel **b**). Each pie chart (panels **c** and **d**) corresponds to one category of the gene pairs specified in the title of the given chart, which indicates the proportion of gene pairs not shared (independent) with other categories (black) and the proportion shared with a given category color-coded according to the legend at the bottom of the figure. It is shown both among all gene pairs (panel **c**) and specifically among synthetic lethal ones (panel **d**). Panels **e)** is the same as figures 2c, but the synthetic lethal gene pairs belonging to the four categories color-coded according to the legend are distinguished from the other synthetic lethal gene pairs, which are not assigned to any of the four categories (grey color). The vertical axis in panel **e)** indicates the number of synthetic lethal gene pairs with genomic distance <25 KB, while in panel **f)** it shows the number of synthetic lethal gene pairs, which are closely located in the 3D chromosomal space of the budding yeast (considering HindIII fragments as one KB).

Finally, we aimed to quantify the contribution of these four classes in the 2D and 3D attraction of synthetic lethal gene pairs. We observed that only 16/161 (9.9%) of gene pairs among the closely located SL pairs (genomic distance < 25 KB) belong to one of these four classes (Figure 4e), which accounts only for 16/97.6 (16.4%) of the difference between the observed and expected closely located SL gene pairs. Hence, even if we remove all the SL gene pairs of these four classes, the enrichment of closely located SL gene pairs remains significant (observed: 145, expected: 63.4, *χ*^2^=105.02 and p-value < 10^−23^). Similarly, we observed only 64/407 (15.7%) of gene pairs among the 3D proximate SL pairs belong to one of these four classes (Figure 4f), which accounts for 64/176.1 (36.3%) of the difference between observed and expected 3D proximate SL gene pairs. Hence, even if we remove all the gene pairs of these four classes, the enrichment of 3D proximate SL gene pairs remains significant (observed: 343, expected: 230.9, *χ*^2^=54.4 and p-value < 10^−12^). Therefore, the enrichment of synthetic lethal gene pairs among the same-pathway, co-expressed, PPI interaction pairs and paralogs, does not sufficiently explain the 2D and 3D attraction of SL gene pairs.

## Discussion

It is well known that the arrangement of genes on chromosomes both in prokaryotes and in eukaryotes is not random, but rather has been shaped under the influence of selective forces over the course of evolution (Adams et al. 1992; Trowsdale 2002; Hurst et al. 2004). Synthetic lethal interactions profoundly impact cellular survival and so it is not unexpected that the relative genomic arrangement of SL gene pairs reveals special evolutionary patterns. Previously, using a computational approach we observed that synthetic lethal (metabolic) gene pairs on bacterial genomes tend to be located further away (“repulsed”) from each other (Hosseini & Wagner 2018). Although ample experimentally verified gene interaction data for *E. coli* has been generated using SGA method (Babu et al. 2011, 2014; Kumar et al. 2016; Gagarinova et al. 2016) the technology suffers from linkage suppression, which prohibits reliable identification of genetic interactions in closely located loci (Typas et al. 2008), and hence the current experimental data in *E. coli* cannot be used to support or refute our previous computational observation.

Fortunately, the linkage suppression problem does not prohibit such an investigation in yeast, because there are orthogonal confirmatory experimental systems such as tetrad analysis, random sporing and spot dilution assays available for yeast, which can minimize the effect of this potential technical artefact by confirming the true positive hits from the pool of candidate synthetic lethal interactions determined by SGA or other high-throughput methods (see methods). This collection of reliably conservative experimental SL data in yeast encouraged us to check the generalizability of the previous computational observation beyond the prokaryotic species. Jacques Monod assertion on biological unity, “*anything found to be true of E. coli must also be true of Elephants*” (Friedmann 2004), resonates well with this generalization purpose, but it needs to be taken with a grain of salt, at least because exceptions pervade biological rules. As a prominent example, the operon organization, an *E. coli* based discovery that earned Jacques Monod the Nobel prize, has rarely been observed in eukaryotes.

More importantly, the evolutionary forces operating on prokaryotic and eukaryotic genomes in some respects are fundamentally different. For example, whereas there is ample evidence attesting to the importance of large-scale gene deletions on the evolution of prokaryotic genomes (Mira et al. 2001; Kunin & Ouzounis 2003; Nilsson et al. 2005; Lee et al. 2012; Koskiniemi et al. 2012; Albalat & Cañestro 2016; Sung et al. 2016), such evidence for eukaryotic ones is rare. Therefore, if the repulsion of synthetic lethal gene pairs is a signature of adaptation to pervasive deletional events as was shown in the previous study (Hosseini & Wagner 2018), it must remain as a particular characteristic of the prokaryotic genomes. Our results indeed revealed evidence against the repulsion of synthetic lethal gene pairs in eukaryotes. Interestingly, we observed even an opposite trend, namely the enrichment of closely located synthetic lethal gene pairs (i.e. attraction) both in 2D and 3D chromosomal space of the *S. cerevisiae*.

To gain mechanistic insights into the origin of the genomic attraction of SL genes in yeast, we identified four relatively independent classes of gene pairs, in which SL interactions are highly enriched: i) gene pairs operating on the same pathways, ii) co-expressed genes, iii) the gene pairs whose protein products physically interact and iv) the paralogs. We observed that unlike the other three classes of genes, the gene pairs whose proteins physically interact are not enriched among the 2D proximate genes. However, all four types of genes were significantly enriched among the 3D proximate ones. Our emphasis on the 3D organization of gene pairs is particularly important, without which we would have neglected, for example, the importance of protein-protein interactions in the chromosomal organization of synthetic lethal genes. This emphasizes the importance of the 3D arrangement of genes towards establishing an evolutionary theory of genome organization in eukaryotes (Hurst et al. 2004). Importantly, we also observed a considerable overlap between the SL gene pairs in the first two classes with the third one, which further underscores the importance of the protein-protein interactions among SL gene pairs that has also been highlighted recently in the human cancer cell lines (Lord et al. 2020).

Although our attempt to dissect the aforementioned four classes of gene pairs is insightful on its own, we concluded that the enrichment of 2D and 3D proximate synthetic lethal gene pairs cannot be fully explained by them, and so its evolutionary origin still remains as an open question. In other words, the proximity of synthetic lethal gene pairs in yeast cannot be fully explained by functional relevance, co-expression, protein-protein interaction or gene duplication events. To understand its origin, we might either need to characterize additional types of gene pairs enriched among SL genes or attribute these patterns directly to natural selection, in which case the next emerging question to be addressed would be what are the direct evolutionary advantages of the 2D and 3D proximity of synthetic lethal genes for a eukaryotic species like the budding yeast?

## Acknowledgements

This work was supported by Open Targets [grant number OT2062 to S-R.H]; and EMBL-EBI [to E.P and B.W].

## Data availability

The data underlying this article are available in the article and in its online supplementary material.

## Supplementary Information

Tables S1-S8 are available at: https://drive.google.com/drive/folders/1D7IUXrDB_noHumGqZj7GbPUCdEiMvo9D?usp=sharing

## Notes

https://drive.google.com/drive/folders/1D7IUXrDB_noHumGqZj7GbPUCdEiMvo9D?usp=sharing

### Competing Interest Statement

The authors have declared no competing interest.

### Summary of Updates

We removed the analyses pertaining to the E. coli and the human synthetic lethality as there were technical problems associated with their data. Instead, in the new version, we focused entirely on the yeast and enhanced the previous study by including additional analyses of the 3D chromosomal arrangement of the synthetic lethal gene pairs in the yeast genome. Moreover, we have provided a more comprehensive picture, by putting synthetic lethality in the context of other important processes such as co-functionality, co-expression, protein-protein interactions and gene duplication. Previous figures 2-4 are replaced with three new figures and the method section is more elaborate in the new revision.

## References

Adams RLP et al. 1992. The arrangement of genes. In: The Biochemistry of the Nucleic Acids. Springer Netherlands pp. 317–338. doi: 10.1007/978-94-011-2290-0_8.

Albalat R, Cañestro C. 2016. Evolution by gene loss. Nat. Rev. Genet. 17:379–391. doi: 10.1038/nrg.2016.39.

Babu M et al. 2011. Genetic interaction maps in Escherichia coli reveal functional crosstalk among cell envelope biogenesis pathways. PLoS Genet. 7. doi: 10.1371/journal.pgen.1002377.

Babu M et al. 2014. Quantitative Genome-Wide Genetic Interaction Screens Reveal Global Epistatic Relationships of Protein Complexes in Escherichia coli. PLoS Genet. 10. doi: 10.1371/journal.pgen.1004120.

Bellay J et al. 2011. Putting genetic interactions in context through a global modular decomposition. Genome Res. 21:1375–1387. doi: 10.1101/gr.117176.110.

Benstead-Hume G et al. 2019. Predicting synthetic lethal interactions using conserved patterns in protein interaction networks. PLoS Comput. Biol. 15. doi: 10.1371/journal.pcbi.1006888.

Cohen BA, Mitra RD, Hughes JD, Church GM. 2000. A computational analysis of whole-genome expression data reveals chromosomal domains of gene expression. Nat. Genet. 26:183–186. doi: 10.1038/79896.

Costanzo M et al. 2016. A global genetic interaction network maps a wiring diagram of cellular function. Science (80-.). 353:42–65. doi: 10.1126/science.aaf1420.

Costanzo M et al. 2010. The genetic landscape of a cell. Science (80-.). 327:425–431. doi: 10.1126/science.1180823.

Dan Tenenbaum. 2018. KEGGREST: Client-side REST access to KEGG. R Packag. version 1.18.1. https://bioconductor.org/packages/release/bioc/html/KEGGREST.html.

Dede M, McLaughlin M, Kim E, Hart T. 2020. Multiplex enCas12a screens show functional buffering by paralogs is systematically absent from genome-wide CRISPR/Cas9 knockout screens. bioRxiv. 2020.05.18.102764. doi: 10.1101/2020.05.18.102764.

Dixon SJ et al. 2008. Significant conservation of synthetic lethal genetic interaction networks between distantly related eukaryotes. Proc. Natl. Acad. Sci. U. S. A. 105:16653–16658. doi: 10.1073/pnas.0806261105.

Duan Z et al. 2010. A three-dimensional model of the yeast genome. Nature. 465:363–367. doi: 10.1038/nature08973.

Friedman R, Hughes AL. 2001. Gene duplication and the structure of eukaryotic genomes [1]. Genome Res. 11:373–381. doi: 10.1101/gr.155801.

Gagarinova A et al. 2016. Systematic Genetic Screens Reveal the Dynamic Global Functional Organization of the Bacterial Translation Machinery. Cell Rep. 17:904–916. doi: 10.1016/j.celrep.2016.09.040.

Hibbs MA et al. 2007. Exploring the functional landscape of gene expression: directed search of large microarray compendia | Bioinformatics | Oxford Academic. Bioinformatics. 23:2692–2699. https://academic.oup.com/bioinformatics/article/23/20/2692/229926 (Accessed June 20, 2020).

Hosseini S-R, Wagner A. 2018. Genomic organization underlying deletional robustness in bacterial metabolic systems. Proc. Natl. Acad. Sci. U. S. A. 201717243. doi: 10.1073/pnas.1717243115.

Hurst LD, Pál C, Lercher MJ. 2004. The evolutionary dynamics of eukaryotic gene order. Nat. Rev. Genet. 5:299–310. doi: 10.1038/nrg1319.

De Kegel B, Ryan CJ. 2019. Paralog buffering contributes to the variable essentiality of genes in cancer cell lines Dudley, AM, editor. PLOS Genet. 15:e1008466. doi: 10.1371/journal.pgen.1008466.

Kelley R, Ideker T. 2005. Systematic interpretation of genetic interactions using protein networks. Nat. Biotechnol. 23:561–566. doi: 10.1038/nbt1096.

Koskiniemi S, Sun S, Berg OG, Andersson DI, Boxer D. 2012. Selection-Driven Gene Loss in Bacteria Casadesús, J, editor. PLoS Genet. 8:e1002787. doi: 10.1371/journal.pgen.1002787.

Kumar A et al. 2016. Conditional Epistatic Interaction Maps Reveal Global Functional Rewiring of Genome Integrity Pathways in Escherichia coli. Cell Rep. 14:648–661. doi: 10.1016/j.celrep.2015.12.060.

Kunin V, Ouzounis CA. 2003. The Balance of Driving Forces During Genome Evolution in Prokaryotes. Genome Res. 13:1589–1594. doi: 10.1101/gr.1092603.

Lee M-C, Marx CJ, Lenski R, Sivam D, Lidstrom M. 2012. Repeated, Selection-Driven Genome Reduction of Accessory Genes in Experimental Populations Moran, NA, editor. PLoS Genet. 8:e1002651. doi: 10.1371/journal.pgen.1002651.

Lord CJ, Quinn N, Ryan CJ. 2020. Integrative analysis of large-scale loss-of-function screens identifies robust cancer-associated genetic interactions. Elife. 9:1–37. doi: 10.7554/eLife.58925.

Mira A, Ochman H, Moran NA. 2001. Deletional bias and the evolution of bacterial genomes. Trends Genet. 17:589–96. http://www.ncbi.nlm.nih.gov/pubmed/11585665 (Accessed February 4, 2014).

Nijman SMB. 2011. Synthetic lethality: General principles, utility and detection using genetic screens in human cells. FEBS Lett. 585:1–6. doi: 10.1016/j.febslet.2010.11.024.

Nilsson AI et al. 2005. Bacterial genome size reduction by experimental evolution. Proc. Natl. Acad. Sci. U. S. A. 102:12112–6. doi: 10.1073/pnas.0503654102.

Oughtred R et al. 2018. The BioGRID interaction database: 2019 update. Nucleic Acids Res. 47:529–541. doi: 10.1093/nar/gky1079.

Pan X et al. 2007. dSLAM analysis of genome-wide genetic interactions in Saccharomyces cerevisiae. Methods. 41:206–221. doi: 10.1016/j.ymeth.2006.07.033.

Qian W, Zhang J. 2014. Genomic evidence for adaptation by gene duplication. Genome Res. 24:1356–1362. doi: 10.1101/gr.172098.114.

Simonis M et al. 2006. Nuclear organization of active and inactive chromatin domains uncovered by chromosome conformation capture-on-chip (4C). Nat. Genet. 38:1348–1354. doi: 10.1038/ng1896.

Srivas R et al. 2016. A Network of Conserved Synthetic Lethal Interactions for Exploration of Precision Cancer Therapy. Mol. Cell. 63:514–525. doi: 10.1016/j.molcel.2016.06.022.

Sung W et al. 2016. Evolution of the Insertion-Deletion Mutation Rate Across the Tree of Life. G3&amp;#58; Genes|Genomes|Genetics. 6:2583–2591. doi: 10.1534/g3.116.030890.

Tong AHY et al. 2001. Systematic genetic analysis with ordered arrays of yeast deletion mutants. Science (80-.). 294:2364–2368. doi: 10.1126/science.1065810.

Trowsdale J. 2002. The gentle art of gene arrangement: The meaning of gene clusters. Genome Biol. 3:comment2002.1. doi: 10.1186/gb-2002-3-3-comment2002.

Typas A et al. 2008. High-throughput, quantitative analyses of genetic interactions in E. coli. Nat. Methods. 5:781–787. doi: 10.1038/nmeth.1240.

Wong SL et al. 2004. Combining biological networks to predict genetic interactions. Proc. Natl. Acad. Sci. U. S. A. 101:15682–15687. doi: 10.1073/pnas.0406614101.

